# Hepatocyte-Specific Hepatocyte Nuclear Factor 4 alpha (HNF4α) Deletion Decreases Resting Energy Expenditure By Disrupting Lipid and Carbohydrate Homeostasis

**DOI:** 10.1101/401802

**Authors:** Ian Huck, E. Matthew Morris, John Thyfault, Udayan Apte

**Affiliations:** Department of Pharmacology, Toxicology and Therapeutics, University of Kansas Medical Center, Kansas City, KS; Department of Molecular and Integrative Physiology, University of Kansas Medical Center, Kansas City, KS; Research Service, Kansas City VA Medical Center

## Abstract

Hepatocyte Nuclear Factor 4 alpha (HNF4α) is required for hepatocyte differentiation and regulates expression of genes involved in lipid and carbohydrate metabolism including those that control VLDL secretion and gluconeogenesis. Whereas previous studies have focused on specific genes regulated by HNF4α in metabolism, its overall role in whole body energy utilization has not been studied. In this study, we used indirect calorimetry to determine the effect of hepatocyte-specific HNF4α deletion (HNF4α-KO) in mice on whole body energy expenditure (EE) and substrate utilization in fed, fasted, and high fat diet (HFD) conditions. HNF4α-KO had reduced resting EE during fed conditions and higher rates of carbohydrate oxidation with fasting. HNF4α-KO mice exhibited decreased body mass caused by fat mass depletion despite no change in energy intake and evidence of positive energy balance. HNF4α-KO mice were able to upregulate lipid oxidation during HFD suggesting that their metabolic flexibility was intact. However, only hepatocyte specific HNF4α-KO mice exhibited significant reduction in basal metabolic rate and spontaneous activity during HFD. Consistent with previous studies, hepatic gene expression in HNF4α-KO supports decreased gluconeogenesis and decreased VLDL export and hepatic β-oxidation in HNF4α-KO livers across all feeding conditions. Together, our data suggest deletion of hepatic HNF4α increases dependence on dietary carbohydrates and endogenous lipids for energy during fed and fasted conditions by inhibiting hepatic gluconeogenesis, hepatic lipid export, and intestinal lipid absorption resulting in decreased whole body energy expenditure. These data clarify the role of hepatic HNF4α on systemic metabolism and energy homeostasis.

## Introduction

Hepatocyte Nuclear Factor 4 alpha (HNF4α) is a nuclear receptor expressed in hepatocytes and is required for hepatocyte differentiation during embryonic development (4, 19), establishing the hepatocyte transcription factor network (12), and maintaining hepatocyte-specific gene expression (17). Important hepatocyte functions, including xenobiotic metabolism, carbohydrate metabolism, fatty acid metabolism, bile acid synthesis, blood coagulation and ureagenesis (6) are putatively regulated by HNF4α. Additionally, recent studies from our laboratory and others have revealed a novel role of HNF4α in inhibition of cell proliferation and tumor suppression; while deletion of HNF4α in adult mice results in spontaneous proliferation (3, 27) and promotes carcinogen-initiated hepatocellular carcinoma (26).

The role of HNF4α in hepatic lipid and carbohydrate metabolism has been well recognized. Hepatocyte-specific HNF4α deletion either postnatally, or in adult mice, results in steatosis and depletion of hepatic glycogen (3, 7, 26). Hepatic steatosis in HNF4α-KO mice has been attributed to a reduced capacity for exporting lipids through VLDL secretion as a consequence of reduced MTTP and ApoB expression, increased hepatic uptake of fatty acids via increased CD36 expression, and decreased fatty acid oxidation due to decreased CPT1a expression (7, 15, 32). This results in decreased serum triglycerides, decreased serum free fatty acids and decreased serum cholesterol (7, 15, 32). HNF4α differentially regulates lipid metabolism in fed and fasted states by repressing PPARα during the fed state and co-activating with PPARα during fasting to induce β-oxidation genes like CPT1a (15). Thus, alterations in these pathways following HNF4α knockout would likely alter systemic lipid metabolism but this has not been examined.

In relation to glucose homeostasis, HNF4α KO mice likely display low hepatic glycogen and low circulating glucose due to both energetic and transcriptional alterations. HNF4α is induced during fasting (15, 29, 33) and activates expression of gluconeogenic genes, G6Pase and PEPCK, through co-activation with PGC1a (21, 33). Furthermore, blunted fatty acid oxidation is known to reduce ATP necessary to fuel gluconeogenesis which could occur due to decreased CPT1a expression in HNF4α-KO hepatocytes

These previous investigations have focused on the role of HNF4α in metabolism by examining hepatic gene expression, histology and serum metabolites. Despite its powerful effect on hepatic glucose and lipid metabolism, the exact role of hepatic HNF4α on systemic energy metabolism and substrate utilization patterns has not been examined. Here we used metabolic stressors of fasting and high fat diet (HFD) feeding combined with indirect calorimetry to examine how targeted hepatic deletion of HNF4α in adult mice effects systemic energy metabolism.

## Materials and Methods

### Animal Care and Tissue Preparation

All animal studies were approved by and performed in accordance with the Institutional Animal Care and Use Committee (IACUC) at the University of Kansas Medical Center. All mice were housed in temperature-controlled conditions (23°C) with a 07:00-21:00 h light phase and 21:00 to 07:00 h dark phase. Two to three-month-old male homozygous HNF4α-floxed mice on C57BL/6J background were injected intraperitoneally with AAV8-TBG-eGFP or AAV8-TBG-CRE resulting in wild-type (WT) and hepatocyte-specific deletion of HNF4α (HNF4α-KO). These vectors were purchased from Penn Vector Core (Philadelphia, PA) and injected as previously described (31). Liver was harvested, weighed and flash frozen in liquid nitrogen at time of euthanasia.

To study the effects of HNF4α deletion in fed and fasted conditions (Figs. 1 and 3, Table 2), HNF4α-floxed littermates were injected with AAV8 to generate WT (n=4) and KO (n=4) groups and were housed individually for one week before entering metabolic cages. While in the cages mice were given *ad libitum* access to normal chow (PicoLab Rodent Diet 20 #5053, LabDiet, St. Louis, MO) for 6.5 days. Chow was removed, and mice were subjected to a 12-hour fasting challenge beginning at the start of the dark cycle. Chow was returned to the cages and mice were given 48 hours of *ad libitum* access to chow before euthanasia and tissue collection.

**Figure 1:**
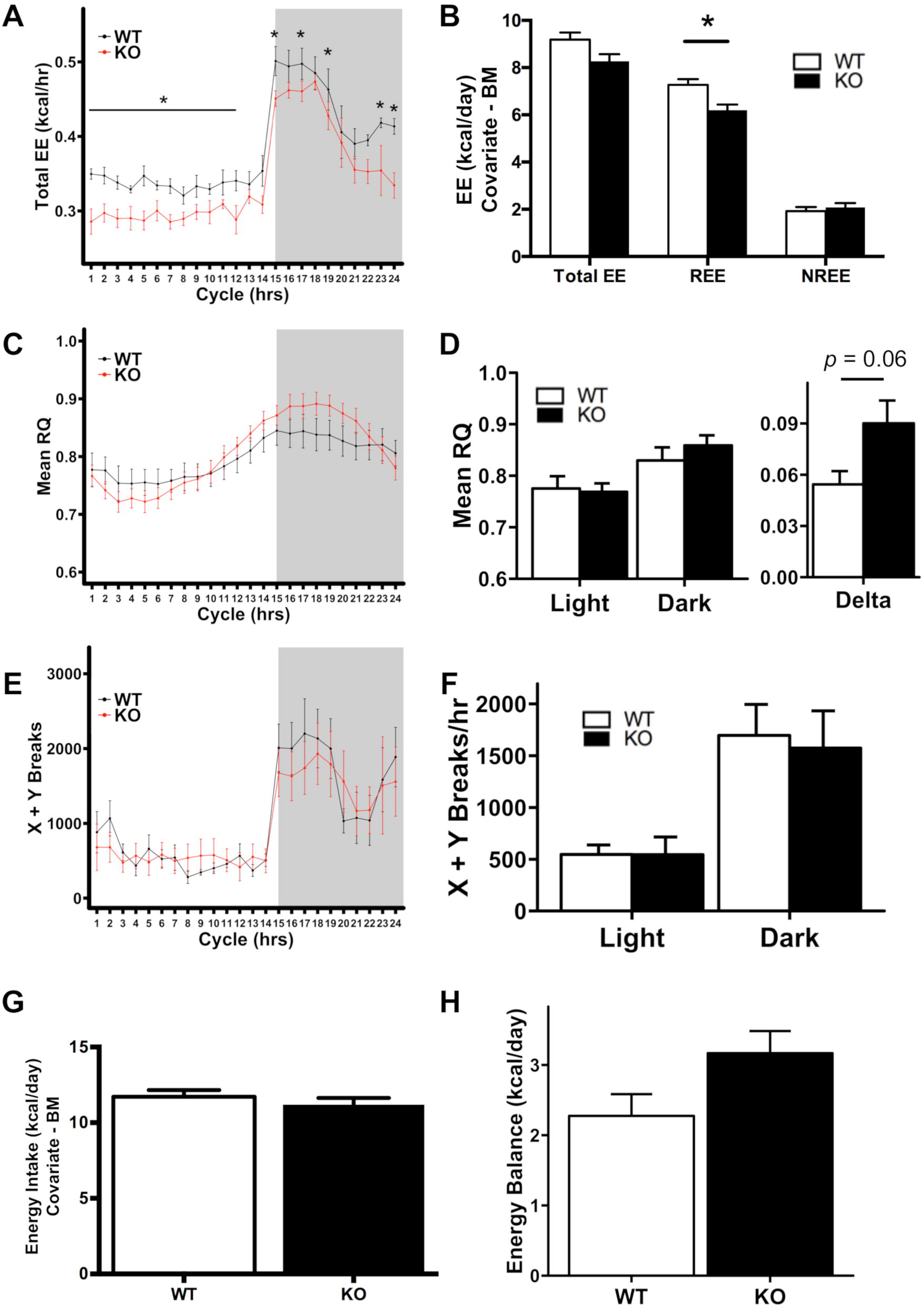
Hepatocyte-specific HNF4α-KO mice exhibit decreased resting energy expenditure during fed conditions. Indirect calorimetry performed on WT and hepatocyte-specific HNF4α-KO mice during 4.5 days of *ad libitum* normal chow feeding. (A) Mean hourly total energy expenditure in WT and HNF4α-KO mice over a 24-hour cycle. Raw values have not been adjusted for body mass differences. (B) Daily total energy expenditure, resting (REE) and non-resting (NREE) energy expenditure in WT and HNF4α-KO mice. Means adjusted with body mass (BM) used as a covariate for energy expenditure. (C) Mean RQ over a 24-hour cycle. (D, Left Panel) Mean RQ summarized by light and dark cycles. (D, Right Panel) Delta depicts the difference in RQ between light and dark cycles. (E) Activity of WT and HNF4α-KO mice expressed as the total X and Y beam breaks for each hour over a 24-hour cycle. (F) Activity of WT and HNF4α-KO mice expressed as the total X and Y beam breaks per hour and summarized per light and dark cycle. (G) Daily energy intake in WT and KO mice with BM used as a covariate. (H) Daily energy balance in WT and KO mice. Gray background indicates dark cycle time point. Values are means ± SEM. Statistical significance determined by one-way ANOVA between WT and KO with * indicating significance with p < 0.05 between WT and HNF4α-KO.

To study the change in energy metabolism and body composition during the process of HNF4α deletion (Fig. 2, Table 3), HNF4α-floxed littermates were housed individually for 5 days before entering metabolic cages. Mice were then acclimatized for 2 days to the indirect calorimetry cage conditions before being injected with AAV8 to generate WT (n=3) and HNF4α-KO (n=4) groups. Mice remained in the cages for 7 days post injection before euthanasia and tissue collection.

**Figure 2:**
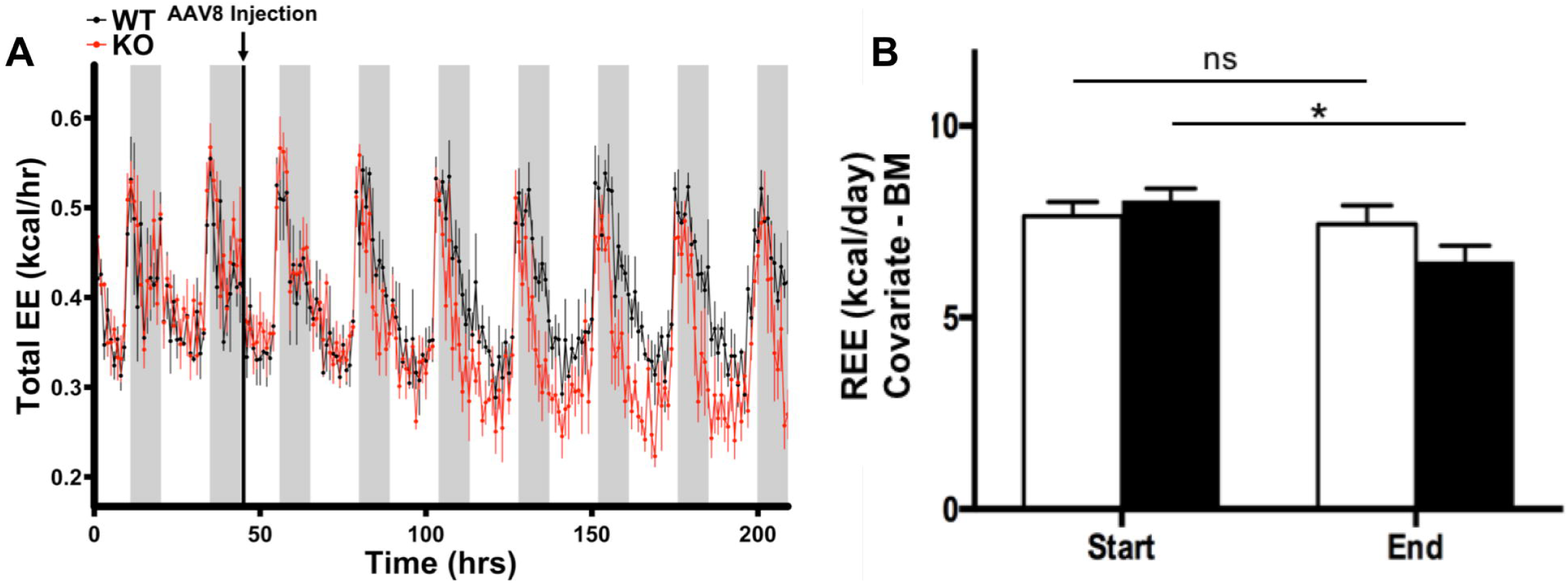
Changes in energy expenditure observed during hepatocyte-specific HNF4α deletion time course. Indirect calorimetry performed on HNF4α-floxed mice before and 7 days after injection with AAV8-TBG-eGFP(WT) or AAV8-TBG-CRE (HNF4α-KO). (A) Mean hourly total energy expenditure (EE) for WT and HNF4α-KO mice before and after AAV8 injection. Raw values have not been adjusted for body mass differences. (B) Resting EE (REE) in WT and HNF4α-KO mice at the start (first 2 days) and end (final 2 days) of the deletion time course. Means adjusted with body mass (BM) used as a covariate for energy expenditure. Gray background indicates dark cycle time point. Values are means ± SEM. Statistical significance determined by one-way ANOVA between WT and KO or between time points with * indicating p < 0.05 between WT and HNF4α-KO.

To study the effects of HNF4α deletion in high fat diet (HFD) conditions, HNF4α-floxed littermates were individually housed, injected with AAV8 to generate WT (n=4) and HNF4α-KO (n=4) groups and fed a control diet containing normal fat content (10% kcal from fat, Cat.# D12110704, Research Diets, Inc., New Brunswick, NJ) for 1 week before entering indirect calorimetry cages. Mice were fed control diet for an additional 3 days in the indirect calorimetry cages before switching to a HFD (60% kcal from fat, catalog# D12492, Research Diets, Inc., New Brunswick, NJ) for 4 days before euthanasia and tissue collection. HFD was replaced every 2 days to prevent spoilage.

### Indirect Calorimetry

Indirect Calorimetry analysis was performed at the University of Kansas Medical Center Metabolic Obesity Phenotyping Facility using a Promethion continuous indirect calorimetry system (Sable Systems International, Las Vegas, NV) as previously described (5). Macros available from the manufacturer were used to calculate total energy expenditure (TEE) based on the Weir equations (28) and respiratory quotient (RQ) by VCO_2_/VO_2_. To calculate resting energy expenditure (REE), the Promethion designed macro pulled out the lowest 30 min period of EE (light cycle) which was then extrapolated to a 24 hour period.. The difference between TEE and REE was used to determine non-resting energy expenditure (NREE). Food was weighed before and after each experiment to determine food consumption. The mass of food consumed was used to determine energy intake (EI) based on the manufacturer provided values for metabolizable energy from each diet type and comparisons were made using body mass as a covariate for EI. Energy Balance (EB) was calculated by subtracting daily total energy expenditure from EI. These calculations were performed as previously described (18). Food hoppers were equipped with a sensor to determine food hopper interaction and manufacturer designed macros used this data to determine hourly feeding bouts. Activity was determined using the XYZ Beambreak Activity Monitor (Sable Systems, Las Vegas, NV) and is expressed as the sum of X and Y beam breaks per time period. Mice were given 2 days to acclimatize to new cages before data was recorded.

### Serum Analysis

Blood glucose was measured with a glucometer using blood collected via retroorbital sinus at time of euthanasia. Serum was isolated and commercially available kits were used to measure β-Hydroxybutyrate (#2440-058, Stanbio), serum triglycerides (#T7532-500, Pointe Scientific) and serum free fatty acids (#MAK044, Sigma-Aldrich).

### Body Composition Analysis

Body composition was performed prior to placement in indirect calorimetry cages and again before euthanasia using the EchoMRI-900 instrument (EchoMRI, Houston, TX). Mice were weighed by hand before entering indirect calorimetry cages, at injection times, before diet changes and before euthanasia. Fat-free mass was calculated by subtracting fat mass from body mass. Percent fat and percent fat free mass was calculated by dividing fat or fat free mass by body mass. Change in fat and fat-free mass was calculated by subtracting the value obtained at the start of the experiment from the value obtained before euthanasia.

### RT-PCR Analysis

RNA from frozen liver was isolated using Trizol method and reverse transcribed to cDNA as previously described (1). Fold change values were calculated in comparison to WT-fed mice using the ddCt method as previously described (14). Table 1 contains primer sequences used in this study.

**Table 1:**
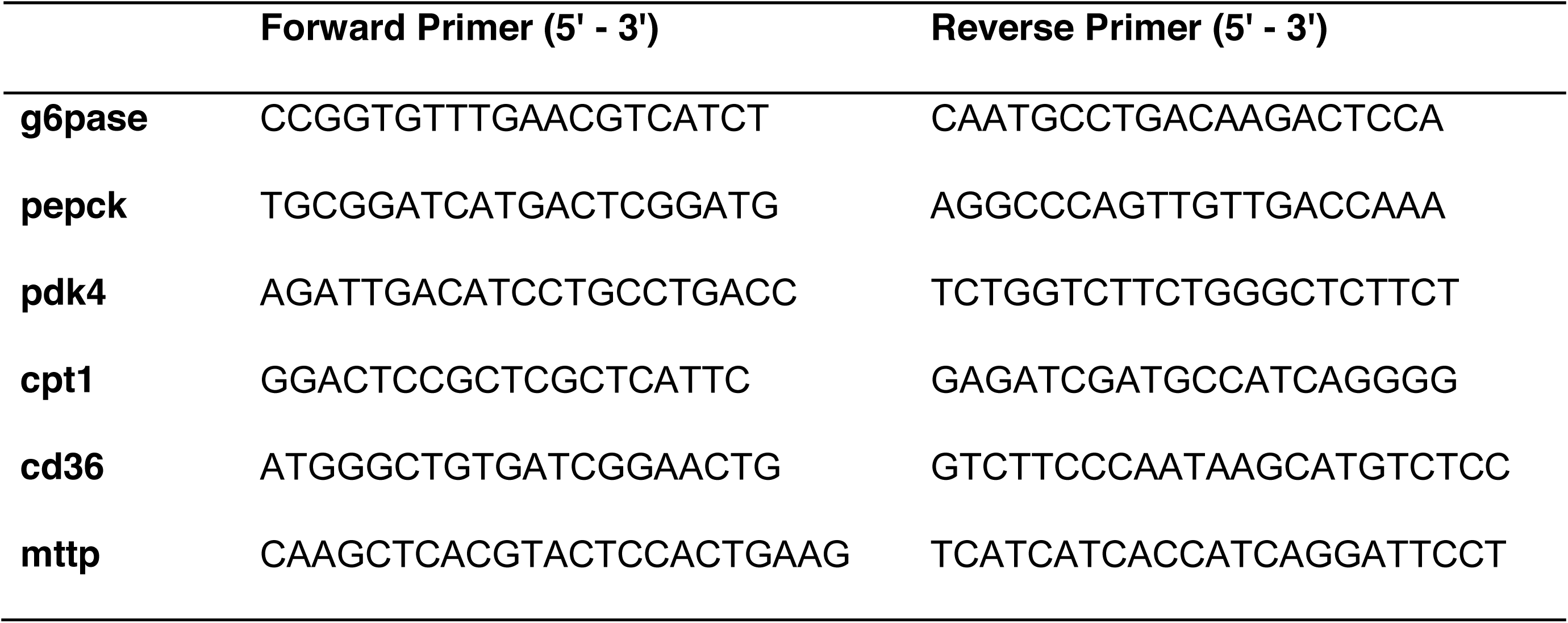
Primer sequences used in this study.

### Statistical Analysis

Statistical analysis was performed using IBM SPSS Statistics Version 25. One Way ANOVA or Student’s t test was applied where indicated with p < 0.05 being considered significant. Analysis of energy expenditure and energy intake with body mass as a covariate was performed as previously described (18) and in accordance with published protocols (24).

## Results

### Hepatocyte-specific HNF4α-KO mice exhibit decreased resting energy expenditure during fed conditions

HNF4α-KO mice on a chow diet exhibited decreased total energy expenditure (TEE) at multiple time points throughout the dark and light cycle (Fig. 1A). TEE and resting energy expenditure (REE) were also significantly lower in HNF4α-KO mice when data was summarized throughout the 24-hour cycle (data not shown). However, energy expenditure is influenced to a significant degree by body mass and multiple methods have been proposed for eliminating the contributing of body mass when comparing energy expenditure in organisms of different size (25). To account for the impact of body mass on energy expenditure, TEE, REE, and non-resting energy expenditure (NREE) were calculated and analyzed with body mass used as a covariate (Fig. 1B). In this experiment, random distribution of littermates into treatment groups resulted in individual differences in body mass, with some HNF4α-KO mice weighing more than their WT littermates. Since these mice also exhibited decreased unadjusted TEE, adjusting means with body mass as a covariate actually underestimated the decrease in EE in HNF4α-KO mice. The correction of EE with body mass as a covariate resulted in the adjusted TEE between WT and HNF4α-KO mice to be similar (Fig. 1B). However, REE values, which were adjusted for body mass, were significantly reduced in HNF4α-KO mice (Fig. 1B). There were no differences in adjusted NREE between genotypes (Fig. 1B). Respiratory quotient (RQ) in HNF4α-KO and WT mice was similar throughout the 24-hour cycle(Fig. 1C). RQ remained similar between WT and HNF4α-KO mice when the data was quantified for the entire light or dark cycle (Fig. 1D, Left Panel).

However, delta RQ, representing the difference in RQ between light and dark cycles, was greater in HNF4α-KO mice and was nearly statistically significant with *p* = 0.06 (Fig. 1D, Right Panel). Activity was measured by total X and Y beam breaks. No significant differences in activity were observed between WT and HNF4α-KO mice at any time point during the 24-hour cycle (Fig. 1E) or when mean hourly beam breaks were summarized per light/dark cycle (Fig. 1F). Daily energy intake (EI) was similar between groups (Fig. 1G). Daily energy balance was similar between groups (Fig. 1H). In summary, on a chow diet the hepatocyte-specific HNF4α-KO mice exhibited decreased resting energy expenditure. This occurred with only minor differences in substrate utilization and no differences in spontaneous activity or energy intake between WT and HNF4α-KO mice. As mentioned previously, individual differences in body mass existed between mice, but there were no significant differences in mean body mass between WT and HNF4α-KO groups (Table 2). There were no significant differences in food consumption (Table 2). HNF4α-KO mice had significantly less fat mass at the end of the experiment and lost a significant amount of fat mass over the 9 days of analysis compared to WT mice (Table 2). Fat-free mass was significantly greater in HNF4α-KO mice (Table 2). HNF4α-KO mice had significantly lower blood glucose compared to WT mice (Table 2).

**Table 2:** Body composition, food consumption and blood glucose data for WT and HNF4α-KO mice fed normal chow. Body mass, fat mass, and blood glucose were measured immediately before euthanasia. Food consumption values depict the mass of normal chow consumed during the entire feeding period and are presented as absolute values. Change in fat mass was calculated for each mouse by subtracting the total fat mass at the start of indirect calorimetry analysis from the fat mass measured immediately before euthanasia. Fat free mass was calculated by subtracting fat mass from body mass using values obtained immediately before euthanasia. Values are means ± SEM. Statistical significance determined by Student’s t-test with * indicating significance with p < 0.05 between WT and HNF4α-KO.

|  | WT | KO |
| --- | --- | --- |
| **Body Mass (g)** | 22.35 ± 0.52 | 23.45 ± 0.21 |
| **Food Consumption (g)** | 24.38 ± 0.36 | 24.13 ± 1.08 |
| **Fat Mass (g)** | 1.83 ± 0.13 | 1.16 ± 0.22* |
| **Change in Fat Mass (g)** | 0.25 ± 0.05 | -0.17 ± 0.16* |
| **Fat Free Mass (g)** | 20.53 ± 0.61 | 22.29 ± 0.15* |
| **Blood Glucose (mg/dL)** | 195.00 ± 12.54 | 102.25 ± 7.82* |

### Changes in energy expenditure observed during hepatocyte-specific HNF4α deletion time course

In the experiments from Figure 1, mice were injected with AAV8 vector and 1 week was allowed for cre recombinase to induce HNF4α deletion before the mice were placed in the indirect system for a 2 day acclimation period. This 9-day time span between viral treatment and data collection produced uncertainty as to whether the reduced REE observed in HNF4α-KO mice was ultimately caused by deletion of HNF4α from hepatocytes, or was a compensatory response to long-term HNF4α deletion. To address this issue, we adjusted the timing of our study so HNF4α-floxed mice were placed into the indirect calorimetry cages for 2 days before being injected with AAV8. EE, RQ and activity was measured for a 7-day time course following injection of AAV8 into mice. If the decreased REE phenotype observed in HNF4α-KO mice was a direct result of HNF4α deletion, we expected REE to be the same between groups prior to AAV8 injection and REE to steadily decrease in mice injected with AAV8-TBG-CRE. As expected, TEE was similar between groups before AAV8 injection. However, TEE began to decrease in HNF4α-floxed mice after AAV8-TBG-CRE injection (Fig 2A), most notably during the light cycle. REE was calculated for WT and HNF4α-KO mice for the initial two days at the start of deletion and the final two days before euthanasia with body mass used as a covariate. REE in WT mice did not change over the experiment time course, but there was a significant decrease in HNF4α-KO over the deletion time course. (Fig. 2B). Although statistically significant differences in REE between WT and HNF4α-KO at both time points were not observed, the trend of decreased REE during HNF4α deletion suggests that the decline in REE would continue in HNF4α-KO mice and become statistically significant over a longer time course, like what was observed in Figure 1. Food consumption was similar between treatments (Table 3). Body mass at time of euthanasia was similar between treatments, but HNF4α-KO mice lost significantly more body mass compared to WT mice from the time of AAV8 injection to euthanasia (Table 3). The decrease in body mass primarily occurred through a decrease in fat mass. Total fat mass was significantly lower in HNF4α-KO mice compared to WT at euthanasia and HNF4α-KO mice lost more total fat mass over the deletion time course compared to WT mice (Table 3). Fat-free mass was similar between treatments (Table 3).

**Table 3:** Body composition data for WT and HNF4α-KO mice in acute deletion study. Body mass and fat mass were measured at the end of the deletion time course immediately before euthanasia. Food consumption values depict the mass of normal chow consumed throughout the entire deletion time course and are presented as absolute values. Change in fat mass was calculated for each mouse by subtracting the total fat mass measured before AAV8 injection from the fat mass measured immediately before euthanasia. Fat free mass was calculated by subtracting fat mass from body mass using the values obtained immediately before euthanasia. Values are means ± SEM. Statistical significance determined by Student’s t-test with * indicating significance with p < 0.05 between WT and HNF4α-KO.

|  | **WT** | **KO** |
| --- | --- | --- |
| **Body Mass (g)** | 26.10 ± 0.83 | 24.78 ± 0.49 |
| **Change in Body Mass (g)** | -0.17 ± 0.38 | -1.40 ± 0.24* |
| **Food Consumption (g)** | 32.37 ± 0.92 | 33.35 ± 3.24 |
| **Fat Mass (g)** | 1.70 ± 0.24 | 1.00 ± 0.13* |
| **Change in Fat Mass (g)** | -0.04 ± 0.03 | -0.59 ± 0.08* |
| **Fat Free Mass (g)** | 24.40 ± 0.69 | 23.77 ± 0.40 |

### HNF4a-KO exhibit elevated RQ during fasting

WT and HNF4α-KO mice were subjected to a 12-hour fasting challenge to determine how this would differently impact EE and substrate utilization. We found no significant differences in TEE between WT and HNF4α-KO mice at any time points throughout the fasting challenge (Fig. 3A). The TEE, REE and NREE summarized throughout the 12 hour fast and adjusted using body mass as a co-variate also did not reveal differences (Fig. 3B). However, we did find pronounced differences in substrate utilization in the HNF4α-KO mice compared to WT. HNF4α-KO mice had a much slower shift in lowering RQ during the fast compared to WT, suggesting a defect in the ability to upregulate the utilization of lipids for energy (Fig. 3C). This metabolic inflexibility phenotype was likely due to a decreased availability of fatty acids from lipolysis as we observed depletion of adipose tissue after HNF4α deletion (Table 2, 3) and found reduced circulating FFA after fasting (Fig. 5A). In addition, reduced fatty acid utilization could also occur because of reduced fat oxidation in the liver due to reduced CPT-1 expression, which is known to be reduced with HNF4α deletion (Fig. 5B). Activity was not different between groups (Fig. 3D).

**Figure 3:**
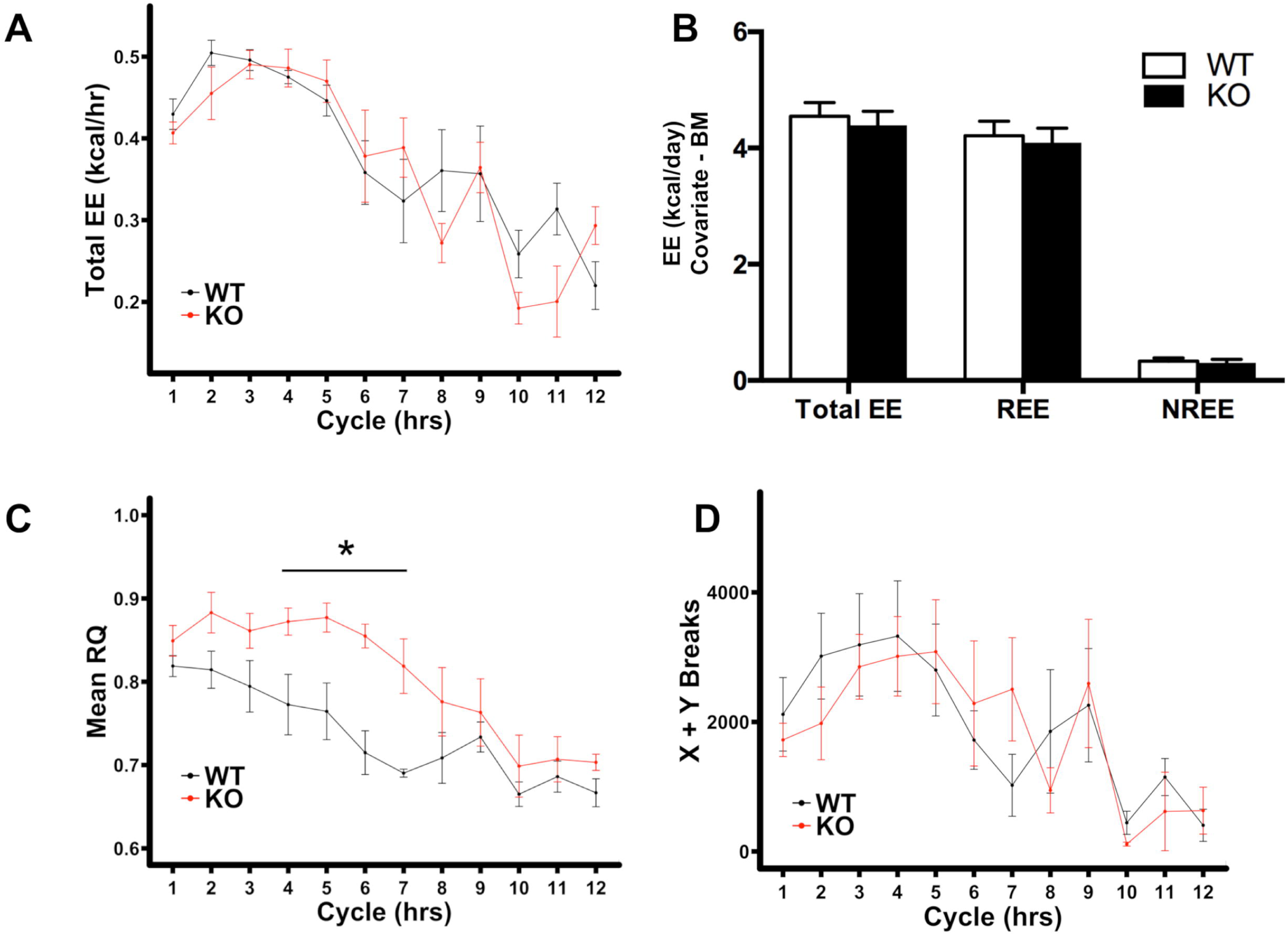
HNF4α-KO mice exhibit elevated RQ during fasting. (A) Mean hourly total energy expenditure (EE) during 12-hour fast. Raw values have not been adjusted for body mass differences. (B) Total EE, resting EE (REE) and non-resting EE (NREE) during 12-hour fast. Means adjusted with body mass (BM) used as a covariate for energy expenditure. (C) Mean hourly RQ during 12-hour fast. (D) Activity during 12-hour fast expressed as total X and Y beam breaks per hour. Values are means ± SEM. Statistical significance determined by one-way ANOVA with * indicating p < 0.05 between WT and HNF4α-KO.

### HNF4α-KO mice exhibit decreased energy expenditure, decreased RQ, decreased activity and loss of body mass during acute high fat diet (HFD) challenge

The metabolic inflexibility displayed by the HNF4α-KO mice to a fasting challenge suggested hepatic HNF4α deletion reduces whole body capacity for lipid utilization. To test the ability of HNF4α to obtain energy from lipids, we monitored energy expenditure, substrate utilization, activity and body composition in WT and HNF4α-KO mice before and after the initiation of a diet containing 60% kcal from fat (HFD). HNF4α-KO mice tended to have lower total EE before HFD feeding. However, total EE was reduced further in HNF4α-KO mice upon HFD feeding (Fig. 4A). Significant reductions in total EE, REE and NREE were observed in HNF4α-KO mice when body mass was used as a covariate (Fig. 4B). Contrary to the metabolic inflexibility observed in HNF4α-KO mice during the fasting challenge, RQ of both groups decreased at the same rate upon initiation of HFD feeding. Further, RQ of HNF4α-KO mice was lower than WT mice at several time points throughout HFD feeding suggesting HNF4α-KO mice had greater capacity for or increased reliance on dietary lipid utilization when consuming a HFD (Fig. 4C,D). Dark cycle activity was significantly reduced in HNF4α-KO mice compared to WT when measured by total X and Y beam breaks (Fig. 4E) as well as feeding bouts (Fig. 4F). Feeding bouts were determined by a manufacturer provided macro, and say more about activity than food consumption, since food intake rate and total food consumption as determined by macros was similar between groups (data not shown). Furthermore, food was weighed by hand at the start and end of the experiment and while HNF4α-KO mice consumed significantly less HFD (Table 4), energy intake was similar between groups when body mass was used as a covariate for food consumption (Fig. 4G). Energy balance was similar between groups and positive in both groups (Fig. 4H). Body mass of WT and HNF4α-KO mice was similar at start of HFD feeding and at euthanasia (Table 4). However, HFD feeding led to significantly different changes in body mass with a 7.7% increase in WT body mass and a 6.2% decrease in HNF4α-KO body mass (Table 4). While comparison of body mass between groups revealed no significant differences, the changes in body mass occurring during HFD feeding were signficicant for both genotypes and were confirmed using the in-cage body mass sensors (Fig. 4I).

**Table 4:**
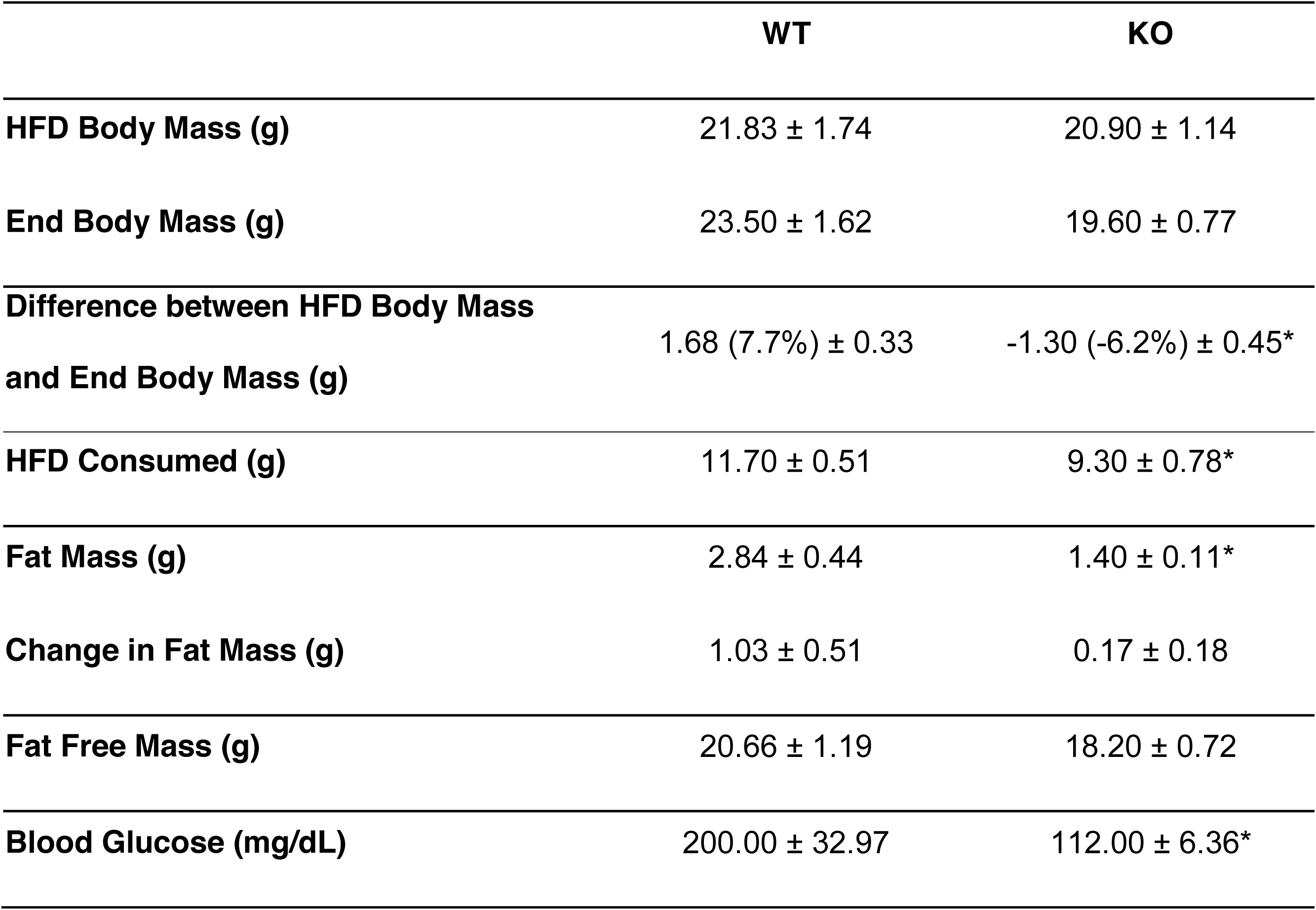
Body composition, food consumption, and blood glucose data for WT and HNF4α-KO mice subjected to acute HFD challenge. HFD body mass was measured when the mice started to feed on HFD. End body mass was measured after 4 days of HFD feeding and immediately before euthanasia. Consumption of HFD represents the total amount consumed throughout the HFD feeding period and are presented as absolute values. Fat mass was measured before euthanasia. Change in fat mass was calculated for each mouse by subtracting the fat mass measured before indirect calorimetry analysis from the fat mass measured before euthanasia. Fat free mass was calculated by subtracting fat mass from body mass using the values obtained immediately before euthanasia. Blood glucose was measured immediately before euthanasia. Values are means ± SEM. Statistical significance determined by Student’s t-test with * indicating significance with p < 0.05 between WT and HNF4α-KO.

|  | WT | KO |
| --- | --- | --- |
| **HFD Body Mass (g)** | 21.83 ± 1.74 | 20.90 ± 1.14 |
| **End Body Mass (g)** | 23.50 ± 1.62 | 19.60 ± 0.77 |
| **Difference between HFD Body Mass**<br>**and End Body Mass (g)** | 1.68 (7.7%) ± 0.33 | -1.30 (-6.2%) ± 0.45* |
| **HFD Consumed (g)** | 11.70 ± 0.51 | 9.30 ± 0.78* |
| **Fat Mass (g)** | 2.84 ± 0.44 | 1.40 ± 0.11* |
| **Change in Fat Mass (g)** | 1.03 ± 0.51 | 0.17 ± 0.18 |
| **Fat Free Mass (g)** | 20.66 ± 1.19 | 18.20 ± 0.72 |
| **Blood Glucose (mg/dL)** | 200.00 ± 32.97 | 112.00 ± 6.36* |

**Figure 4:**
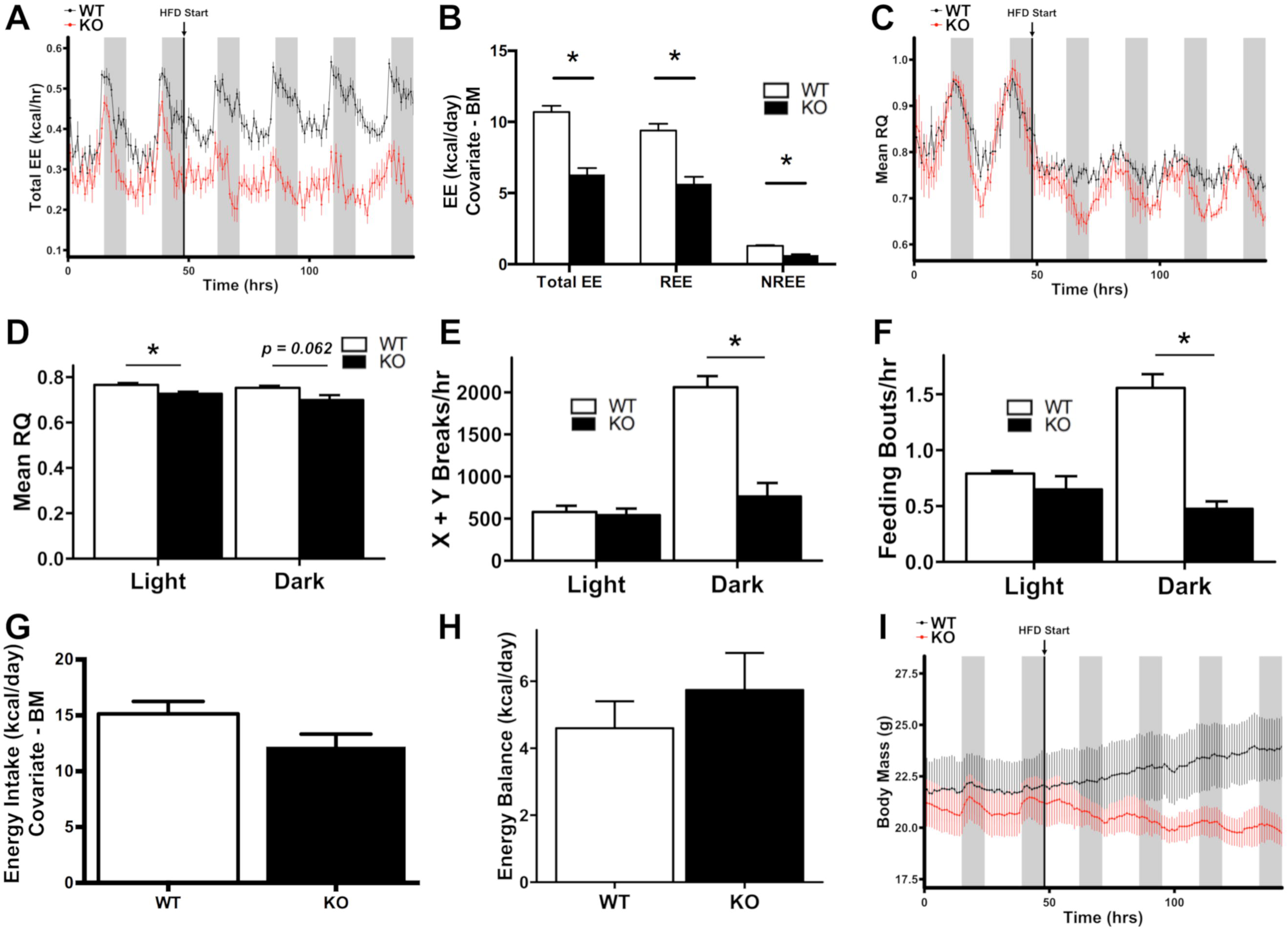
HNF4α-KO mice exhibit decreased energy expenditure, decreased RQ, decreased activity and loss of body mass during acute high fat diet (HFD) challenge. (A) Mean hourly total energy expenditure (EE) during the transition from normal chow to HFD. Raw values have not been adjusted for body mass differences. (B) Total EE, resting EE (REE) and non-resting EE (NREE) in WT and HNF4α-KO mice fed HFD. Means adjusted with body mass (BM) used as a covariate for energy expenditure. (C) Mean hourly RQ during transition from normal chow to HFD. (D) Mean RQ during HFD summarized by light and dark cycles. (E) Activity in HFD fed WT and HNF4α-KO mice during light and dark cycles expressed as the total X and Y beam breaks per hour. (F) Hourly feeding bouts during light and dark cycle in WT and HNF4α-KO fed HFD. (G) Daily energy intake in HFD fed WT and HNF4α-KO mice. Means adjusted with body mass used as a covariate for energy intake. (H) Daily energy balance in HFD fed WT and HNF4α-KO mice. (I) Average body mass in WT and HNF4α-KO mice during transition from normal chow to HFD. Gray background indicates dark cycle time points. Values are means ± SEM. Statistical significance determined by one-way ANOVA with * indicating significance with p < 0.05 between WT and HNF4α-KO.

HNF4α-KO mice contained significantly less fat mass than WT mice after HFD feeding (Table 4). Comparison of the change in fat mass before and after HFD was similar between groups (Table 4). Fat free mass was similar between WT and HNF4α-KO mice (Table 4). Blood glucose was significantly lower in HNF4α-KO mice compared to WT mice, reflective of impaired gluconeogenesis (Table 4).

### Changes in serum lipids and hepatic gene expression in WT and HNF4α-KO mice across feeding conditions

To understand the metabolic phenotype characterized using indirect calorimetry, we measured serum lipids, ketones and hepatic expression of lipid and carbohydrate metabolism genes in WT and HNF4α-KO mice during fed, fasted and HFD conditions. Serum triglycerides and serum free fatty acids were significantly lower in HNF4α-KO animals in all conditions (Fig. 5A). No significant differences in serum ketones were observed between WT and HNF4α-KO mice (Fig. 5A). Gene expression for the rate limiting enzymes in gluconeogenesis (G6Pase, PEPCK) were decreased in HNF4α-KO mice in all conditions as expected given their lower glucose levels (Fig. 5B). PDK4, an enzyme involved in conservation of glucose and upregulation of lipid utilization, was elevated in HNF4α-KO mice in all conditions (Fig. 5B). CPT1, the rate limiting enzyme for fatty acid entry into the mitochondria and subsequent β-oxidation, was suppressed in HNF4α-KO mice in all conditions (Fig. 5C). Expression of the hepatic lipid importer CD36 was elevated in HNF4α-KO mice in all conditions (Fig. 5C). Expression of the apolipoprotein packaging enzyme MTTP was suppressed in HNF4α-KO mice in all conditions (Fig. 5C). Thus, gene expression profiles of the livers from the HNF4α-KO are suggestive of impaired gluconeogenesis and enhanced lipid uptake paired with reduced capacity to oxidize and export lipids.

**Figure 5:**
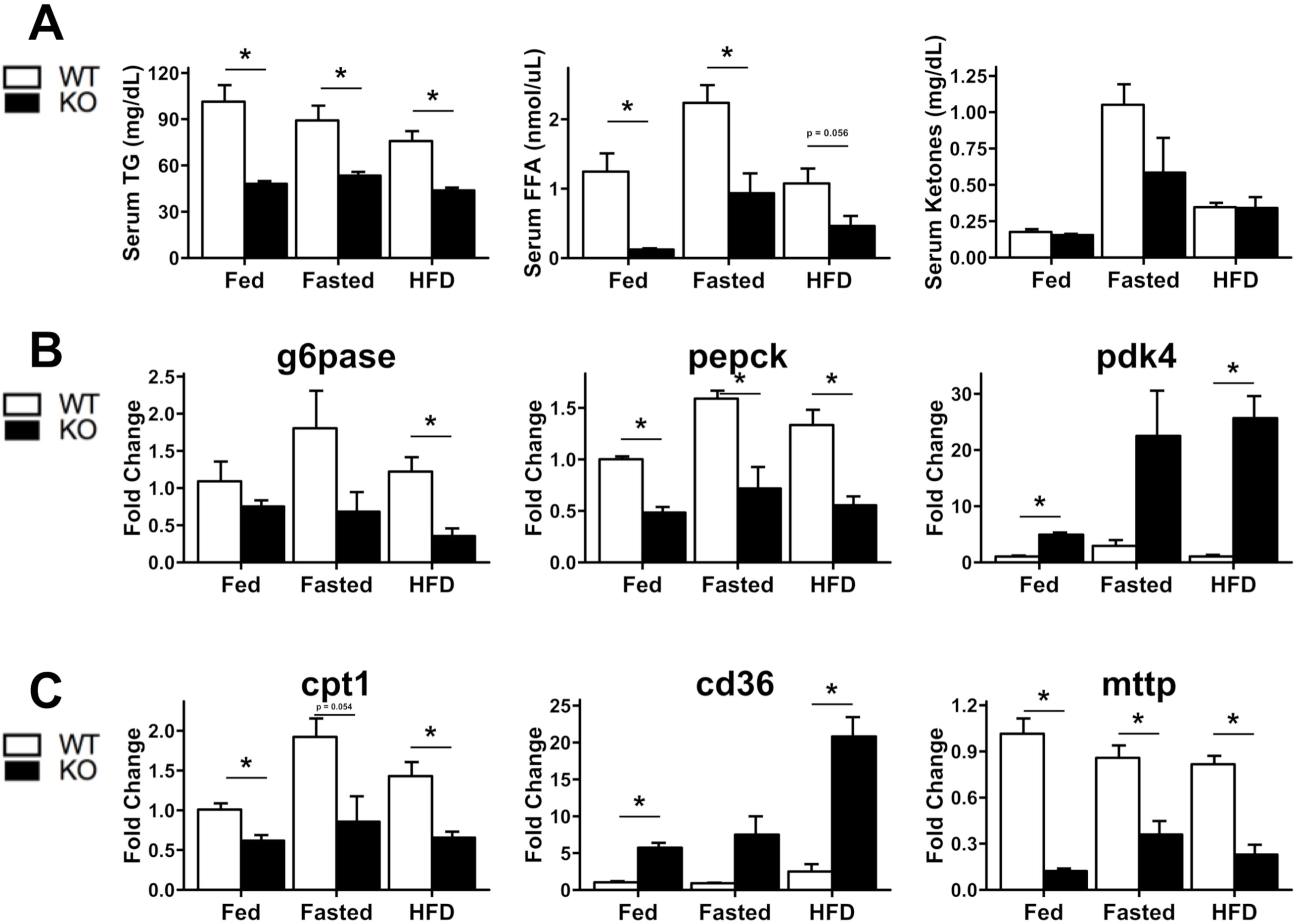
Changes in serum lipids and hepatic gene expression in WT and HNF4α-KO mice across feeding conditions. (A) Serum quantification of (left to right) triglycerides (TG), free fatty acids (FFA) and β-hydroxybutyrate (Ketones). Hepatic expression of genes involved in (B) glucose metabolism and (C) lipid metabolism and trafficking. Gene expression expressed as fold change compared to WT mice fed normal chow. Values are means ± SEM. Statistical significance determined by one-way ANOVA with * indicating significance with p < 0.05 between WT and HNF4α-KO.

## Discussion

The role of HNF4α in hepatocyte differentiation and quiescence is well recognized (4, 12, 13, 17, 19). HNF4α regulates several liver specific functions including lipid and carbohydrate metabolism, but our understanding of the role of HNF4α in metabolism is limited to its role in regulating hepatic gene expression and hepatic nutrient storage. Whether HNF4α mediated regulation of hepatic metabolism results in changes to systemic energetics has not been studied, which was the focus of these studies.

The major finding of this study was that deletion of HNF4α from hepatocytes results in decreased resting energy expenditure during fed conditions. Additionally, hypoglycemia, body mass decrease, and decreased adiposity were observed in HNF4α-KO mice in all feeding conditions. The origin of these phenotypes was directly linked to decreased hepatic HNF4α levels by examining mice during a 7-day time course of hepatic HNF4α deletion. The differences in energy expenditure between WT and HNF4α-KO mice were exaggerated during HFD feeding, with HNF4α-KO mice exhibiting significant decreases in total, resting, and non-resting energy expenditure as well as decreased spontaneous activity.

Reductions in energy expenditure caused by hepatic HNF4α deletion cannot be attributed to decreased energy intake. HNF4α-KO mice exhibited positive energy balance that was similar to WT energy balance during all feeding conditions. Despite exhibiting positive energy balance, HNF4α-KO mice consistently exhibited decreases in body mass which were caused by depletion of fat mass. While it is not clear how HNF4α-KO mice lose body mass while maintaining positive energy balance, it could be explained by poor intestinal lipid absorption due to decreased bile acid production. It is well known that HNF4α regulates genes involved in bile acid synthesis, conjugation and transport (7-9). Cyp7a1, Cyp27a1 and Cyp8b1 are all positively regulated by HNF4α and exhibit decreased expression in HNF4α-KO mice (9). As a result, HNF4α-KO mice synthesize fewer bile acids and excrete fewer bile acids in feces (9). Basal expression of Cyp7a1 requires HNF4α and LRH-1 (11). Like HNF4α, LRH-1 also regulates Cyp8b1 and liver specific deletion of LRH-1 decreases Cyp8b1 expression resulting in decreased bile acid production and poor intestinal lipid absorption (16). It is highly probable that decreased bile acid synthesis in HNF4α-KO mice produces a similar phenotype of reduced intestinal lipid absorption. This would cause HNF4α-KO mice to have increased dependence on endogenous adipose stores for lipid fuel molecules. This is supported by the increased lipolysis observed in HNF4α-KO mice throughout all experiments, including HFD challenge, which was so severe that adipose tissue from epididymal fat pads could not be collected from HNF4α-KO mice. Impaired availability of dietary lipids and dependence on endogenous lipids would also explain the dramatic reductions in HNF4α-KO energy expenditure and activity observed when the majority of dietary caloric intake was available as lipids during the HFD challenge, as well as their resistance to HFD-induced increases in adiposity. Decreased lipid absorption would result in an overestimation of energy intake in HNF4α-KO mice since energy intake was calculated using the mass of food consumed. If the energy actually absorbed by the intestines could be calculated, HNF4α-KO energy balance would likely be negative and reflect the observed loss in body mass.

Decreased availability of dietary lipids for energy production could affect metabolic activity in peripheral lean tissue. Serum triglycerides and free fatty acids were decreased in HNF4α-KO mice in all feeding conditions consistent with previous publications (7, 15, 32). These studies attribute decreased VLDL secretion in HNF4α-KO mice to decreased expression of MTTP and ApoB, which we also observed in all feeding conditions. Decreased hepatic VLDL secretion, combined with depletion of adipose tissue after long term HNF4α deletion, would effectively starve peripheral lean tissue of lipid energy molecules, resulting in decreased energy expenditure.

Additional explanations exist for why we observed decreased energy expenditure after hepatic HNF4α deletion. HNF4α regulates hepatic CPT1a, the enzyme involved in the rate limiting step of fatty acid oxidation, during fed and fasted conditions (15). CPT1a was downregulated in HNF4α-KO mice across all feeding challenges in the current study. It has been estimated that liver accounts for 20% of total EE (20), so decreased capacity for hepatic β-oxidation could result in a detectable reduction in whole body energy expenditure. Decreased hepatic β-oxidation would increase RQ, opposite of what was observed in HNF4α-KO mice. However, this increase in RQ is likely masked the decreased RQ of peripheral tissues which would be forced to metabolize lipids in a hypoglycemic environment.

It has been shown that during cold exposure, activation of HNF4α is required for hepatic synthesis of acyl-carnitines which are used as fuel in brown fat thermogenesis (23). The mice in our study were housed at 24°C. While this is not cold exposure, it could be considered a source of minor thermoregulatory stress (25). If hepatic HNF4α deletion results in reduced acyl-carnitine synthesis, brown adipose tissue in HNF4α-KO mice would be starved of energy molecules required for thermogenesis and therefore be less metabolically active, resulting in decreased energy expenditure.

A previously unknown phenotype of hepatocyte-specific HNF4α deletion was reduction in fat mass. In all experiments with normal chow fed mice, depletion of adipose in HNF4α-KO mice drove a decrease in body mass. Additionally, HNF4α-KO mice were resistant to HFD-induced increases in body mass and adiposity. Increased lipolysis was likely caused by decreased availability of dietary lipids, but could also be caused by increased glucagon signaling in the hypoglycemic HNF4α-KO mice. Long term, this could result in near total lipolysis of adipose therefore preventing a fasting induced increase in serum FFA, such as what was observed in fasted HNF4α-KO mice. White adipose tissue is metabolically active itself and it has been shown to increase energy expenditure in lean tissue via leptin signaling (10). Because HNF4α-KO mice are leaner, leptin signaling is likely lower, which would decrease energy expenditure in lean tissue.

Peripheral tissues of HNF4α-KO mice had decreased access to lipid energy molecules, but were also starved of glucose which could contribute to decreased energy expenditure. In all conditions, HNF4α-KO mice were significantly hypoglycemic. HNF4α is required for PGC1α mediated gluconeogenesis by co-activating PEPCK and G6Pase (21, 33). Moreover, hepatic fat oxidation, which is impaired in HNF4α-KO mice due to decreased CPT1a expression, is necessary for fueling gluconeogenesis. Indeed, PEPCK and G6Pase expression decreased in all feeding conditions following HNF4α deletion. Hepatocyte-specific HNF4α-KO mice exhibit depletion of hepatic glycogen (26) which is likely caused by failed gluconeogenesis. Since liver is the primary gluconeogenic organ and HNF4α is required for hepatic gluconeogenesis, diet must be the exclusive source of carbohydrate for HNF4α-KO mice. Furthermore, if intestinal lipid absorption is impaired in HNF4α-KO mice, dietary carbohydrates would be especially important for energy production. This is supported by the decreases in total energy expenditure in HNF4α-KO mice during the deletion time course which occurred almost exclusively during the light cycle when mice reduce food consumption. Further, when availability of dietary carbohydrates was reduced, such as during HFD challenge, energy expenditure and activity are significantly reduced as HNF4α-KO mice must produce energy from a dwindling supply of endogenous lipid depots along with a liver exhibiting impaired gluconeogenesis.

Substrate utilization data supports the hypothesis that HNF4α-KO mice depend exclusively on diet as a source glucose. During fed conditions, HNF4α-KO mice exhibited nearly significant increased oscillation between light and dark cycle RQ. Close inspection of RQ in Figure 1A suggests a trend of increased dark cycle RQ and decreased light cycle RQ. Decreased light cycle RQ was also observed in HNF4α-KO mice before initiation of HFD feeding (Fig. 4C). Elevated dark cycle RQ in HNF4α-KO mice can be explained by metabolism of dietary glucose consumed during the dark cycle feeding period in an organism with impaired VLDL secretion and exhausted adipose stores. During the light cycle, HNF4α-KO mice stopped feeding and quickly exhausted stored glucose, forcing them to use stored adipose resulting in reduced light cycle RQ.

The most notable difference in substrate utilization between WT and HNF4α-KO mice was observed during the fasting challenge, where HNF4α-KO mice were slow to decrease RQ, suggesting a reluctance for HNF4α-KO mice to use lipids during caloric restriction. However, this was not supported by the HFD challenge, where HNF4α-KO did not have problems using lipids as a fuel source during HFD feeding and actually displayed lower RQ than WT mice during HFD feeding. The dependence on carbohydrates for fuel exhibited by fasted HNF4α-KO mice may not be caused by defective lipolysis or fatty acid oxidation, but rather be a product of deficient hepatic VLDL secretion and exhausted adipose stores following prolonged HNF4α deletion. This would force peripheral tissues to use any remaining carbohydrate for fuel during caloric restriction. As for substrate utilization during the HFD challenge, it is surprising that RQ was lower in HNF4α-KO mice considering dietary lipid absorption was likely reduced. HFD contained very low quantities of carbohydrates and gluconeogenesis was significantly inhibited in HNF4α-KO mice. Thus, options for energy sources in this scenario would be limited to endogenous lipids or residual intestinal lipid absorption. Also, increased ketogenesis could explain the low RQ exhibited by HNF4α-KO mice during HFD feeding (22). Previous studies have shown increased ketosis in HNF4α-KO mice (15), but we observed no differences between WT and HNF4α-KO serum β-hydroxybutyrate. It is possible that ketone production was higher in HNF4α-KO mice during some phases of the dark cycle when RQ was below 0.7 but was not detectable in our samples which were collected during the light cycle. Another explanation could be the decrease in activity of HNF4α-KO mice fed HFD. While HNF4α-KO mice did not consume less energy than WT mice, reduced ambulatory activity during the feeding period translated into fewer dark cycle feeding bouts, which could result in lower RQ.

In conclusion, this study uses indirect calorimetry and targeted hepatic HNF4α deletion in adult mice to address unanswered questions regarding the role of hepatic HNF4α on systemic metabolic homeostasis and energy flux. Hepatocyte-specific HNF4α deletion reduces peripheral availability of energy molecules likely by reducing lipid absorption, hepatic VLDL secretion, and gluconeogenesis resulting in decreased resting energy expenditure. This study demonstrates how changes in hepatic HNF4α, a condition common in disease human liver (2, 30), can significantly impact whole body energy expenditure and availability of energy stores. This data will inform future attempts to target or modulate hepatic HNF4α by demonstrating its role in metabolism outside of the liver. Our findings reveal for the first time the impact of hepatic HNF4α on systemic energy flux and highlight the importance of liver as a central metabolic organ.

